# Deep-sea mussels from a hybrid zone on the Mid-Atlantic Ridge host genetically indistinguishable symbionts

**DOI:** 10.1101/2020.09.11.282681

**Authors:** Merle Ücker, Rebecca Ansorge, Yui Sato, Lizbeth Sayavedra, Corinna Breusing, Nicole Dubilier

## Abstract

The composition and diversity of animal microbiomes is shaped by a variety of factors, many of them interacting, such as host traits, the environment, and biogeography. Hybrid zones, in which the ranges of two host species meet and hybrids are found, provide natural experiments for determining the drivers of microbiome communities, but have not been well studied in marine environments. Here, we analysed the composition of the symbiont community in two deep-sea, *Bathymodiolus* mussel species along their known distribution range at hydrothermal vents on the Mid-Atlantic Ridge, with a focus on the hybrid zone where they interbreed. In-depth metagenomic analyses of the sulphur-oxidising symbionts of 30 mussels from the hybrid zone, at a resolution of single nucleotide polymorphism analyses of ∼2500 orthologous genes, revealed that parental and hybrid mussels have genetically indistinguishable symbionts. While host genetics does not appear to affect symbiont composition in these mussels, geographic location of the mussels on the Mid-Atlantic Ridge explained 45 % of symbiont genetic variability based on redundancy analyses. We hypothesize that geographic structuring of the free-living symbiont population plays a major role in driving the composition of the microbiome in these deep-sea mussels.

## Introduction

The community composition of an animal’s microbiome is the product of multiple interacting factors that include the environment, geography and host genetics (Benson et al., 2010; Davenport, 2016; Rothschild et al., 2018; Spor et al., 2011; Yatsunenko et al., 2012). To which extent host genetics affect microbiome composition is currently a topic of intense debate, in part as high-throughput sequencing is revealing the genetic makeup of host and symbiont populations in ever higher resolution (Di Bella et al., 2013; Ellegren, 2014; Luikart et al., 2003). Animal hybrids are useful for assessing the effects of host genotype on microbiomes (Lim & Bordenstein, 2020). Studies of lab-reared animal hybrids, such as wasps (Brucker & Bordenstein, 2013), fish (Li et al., 2018; Rennison et al., 2019; Sevellec et al., 2019) and mice (Korach-Rechtman et al., 2019; Wang et al., 2015) found that these hosts had different gut microbiota than their parental species, based on sequencing of the microbial 16S rRNA gene. These altered gut microbiomes of hybrids affected the fitness of some hosts, suggesting that microbiomes play an important role in determining species barriers (Brucker & Bordenstein, 2013). Studies on lab-reared hosts cannot, however, fully reflect the environmental conditions animals experience in their natural habitat. Hybrid zones, in which parental species interbreed and produce hybrid offspring, are excellent natural experiments for teasing apart the impact of host genotype, environment and geographic distance on microbiome composition. Yet surprisingly few studies have investigated the microbiota of hybrids from the wild, and these have yielded mixed results. For example, in a hybrid zone of the European house mouse, the gut microbiota of hybrids differed from that of their parental species (Wang et al., 2015). In contrast, in African baboons there were no significant differences between hybrids and their parental species, and gut community composition was best explained by the environment (Grieneisen et al., 2019). To date, such studies, whether on lab-reared animals or those from the wild, have been based on the sequencing of only a few microbial genes, with the vast majority of analyses based on the 16S rRNA gene, or only a variable region of this gene. These studies were therefore limited to determining microbial community composition at the genus level or higher, and could not distinguish closely related species or strains.

Almost nothing is known about the microbial communities of hosts from marine hybrid zones, despite the pervasiveness of such zones in many regions of the oceans. Hydrothermal vents on the Mid-Atlantic Ridge (MAR), an underwater mountain range extending from the Arctic to the Southern Ocean, provide an ideal setting for investigating the microbiomes of hosts in natural hybrid zones. Many of the vents on the MAR are dominated by *Bathymodiolus* mussels that live in a nutritional symbiosis with chemosynthetic bacteria. Two mussel species colonise the northern MAR, *B. azoricus*, which is found at vents from 38°N to 36°N, and *B. puteoserpentis*, which inhabits vents further south from 23°N to 13°N. A hybrid zone between these two host species occurs at the Broken Spur vent field at 29°N on the MAR, where *B. puteoserpentis* co-occurs with hybrids between *B. azoricus* and *B. puteoserpentis* (Breusing et al., 2017; O’Mullan et al., 2001; Won, Hallam, O’Mullan, & Vrijenhoek, 2003).

The symbionts of bathymodiolin mussels are transmitted horizontally from the environment to juvenile mussels, yet each mussel species harbours a highly specific symbiont community (Dubilier et al., 2008; Van Dover et al., 2002; Won, Hallam, O’Mullan, Pan, et al., 2003). This specificity suggests that the genetics of bathymodiolin mussels plays an important role in determining symbiont composition. In this study, we took advantage of the natural hybrid zone of *Bathymodiolus* mussels at the Broken Spur vent field to investigate how host genotype, geographic distance, and the vent environment affect the composition of their sulphur-oxidising (SOX) symbionts. The recent discovery of a high diversity of SOX symbiont strains in *Bathymodiolus* from the MAR, with as many as 16 strains co-occurring in single *Bathymodiolus* mussels (Ansorge et al., 2019; Ikuta et al., 2016; Picazo et al., 2019), made it critical to resolve genetic differences at the strain level of the SOX symbiont community (strain is defined here as suggested by Van Rossum et al., 2020, as subordinate to subspecies, in our study >99 % average nucleotide identity). We achieved this resolution through multilocus phylogeny, genome-wide gene profiling, and single nucleotide polymorphism (SNP)-based population differentiation analyses of 30 *Bathymodiolus* hybrid and parental individuals collected in 1997 and 2001 at the Broken Spur vent field.

## Materials & Methods

A detailed description of samples and methods is available in the Supplementary Information and an overview of the analyses of SOX symbionts used in this study is provided in Supplementary Table S 4. Data files and scripts used for the analyses can be found in the GitHub repository (https://github.com/muecker/Symbionts_in_a_mussel_hybrid_zone).

Broken Spur parental mussels (13 *B. puteoserpentis*) and hybrids (17 F2 – F4 generation hybrids, see supplement) were identified as described previously (Breusing et al., 2016, 2017) (no parental *B. azoricus* were found at Broken Spur). Briefly, mussels were genotyped based on 18 species-diagnostic markers and identified as parental or hybrid mussels using bioinformatic analyses of population structure, admixture and introgression (Supplementary Table S 2). After DNA extraction and sequencing, we assembled metagenomes per mussel individual from Illumina short-read sequences. Metagenome-assembled genomes (MAGs) of the SOX symbionts from each mussel individual were binned (for statistics of symbionts MAGs, see Supplementary Table S 3), representing the consensus of all SOX symbiont strains in each host individual.

To evaluate genetic differences between symbionts from the northern MAR at the level of bacterial subspecies (sensu Van Rossum et al., 2020, here between 97 % and 99 % average nucleotide identity), we used 171 single-copy, gammaproteobacterial marker genes for phylogenomic analysis of the SOX symbiont MAGs and their closest symbiotic, e.g. symbionts of *B. azoricus* from vents north of Broken Spur and *B. puteoserpentis* mussels from vents south of Broken Spur, and free-living relatives (see Supplementary Table S 5). To understand which factors affect symbiont composition on the strain level at the northern MAR, we assessed the influence of geographic distance, host species, vent type (basaltic versus ultramafic rock) and depth on SOX symbiont allele frequencies using redundancy analysis (RDA). We analysed Broken Spur symbiont MAGs at the genome-wide level by comparing their average nucleotide identities (ANI) to resolve differences on the subspecies level. To resolve strain-level differences between SOX symbionts from Broken Spur, we analysed pairwise F_ST_ values based on SNPs in 2496 orthologous genes from Broken Spur SOX symbiont MAGs. To identify genes that differed between hybrid and parental symbiont populations, we analysed the presence/absence and differential abundance of these orthologues, and further investigated pairwise F_ST_ values of all 2496 orthologous genes.

## Results & Discussion

Phylogenomic analysis of 171 single-copy genes revealed the presence of two SOX symbiont subspecies, one specific to *B. azoricus* from the more northern vents Menez Gwen, Lucky Strike and Rainbow, and one specific to *B. puteoserpentis* from the vents further south, Logatchev and Semenov (Figure 1 A, C).

**Figure 1.**
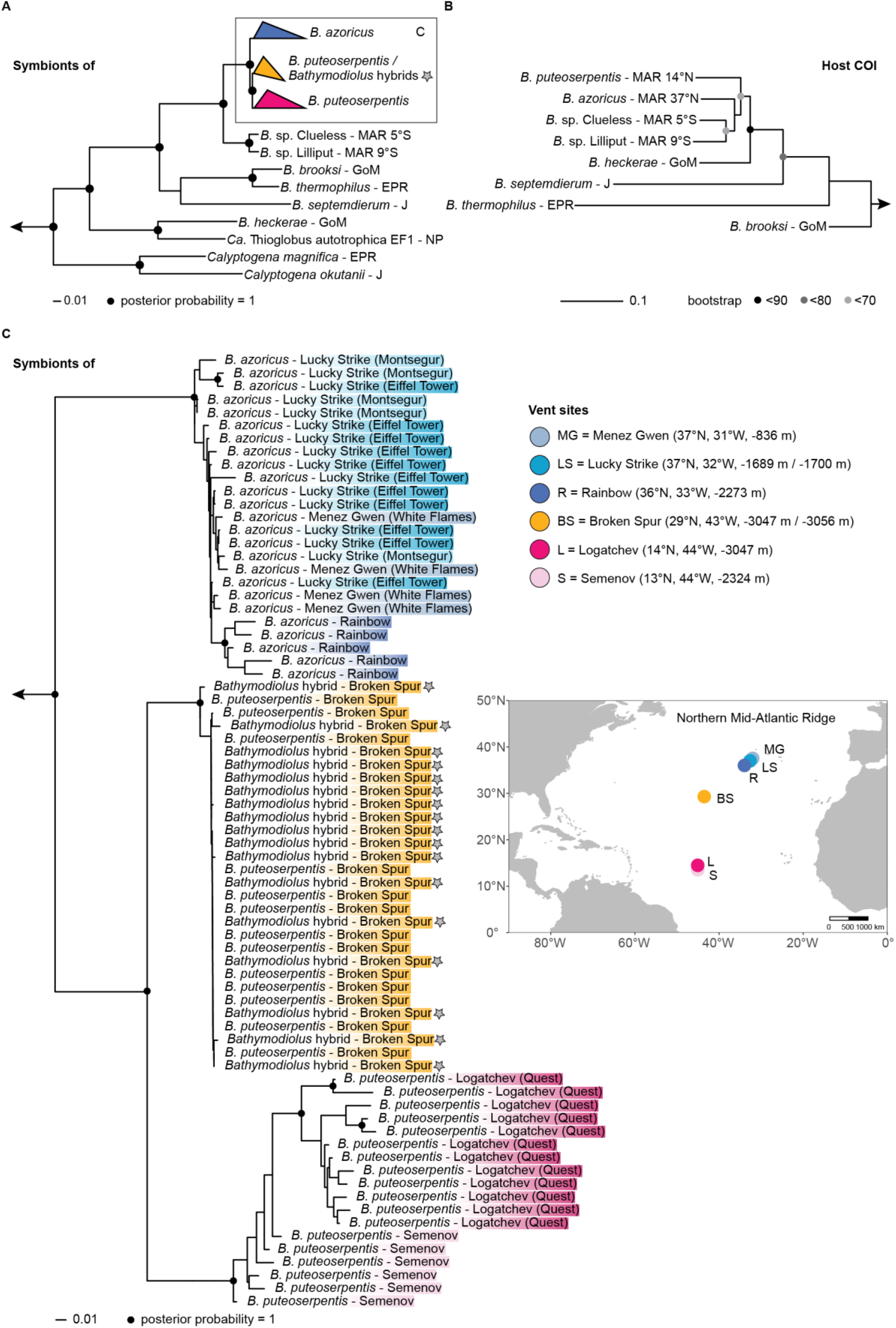
Phylogenetic relationships of *Bathymodiolus* SOX symbionts and their mussel hosts. (A) Overview tree based on 171 single-copy marker genes. The *Bathymodiolus* SOX symbionts from the northern Mid-Atlantic Ridge (blue, yellow and pink) form a clade within the gammaproteobacterial SUP05 clade. *Thiomicrospira* spp. and *Ca*. T. singularis PS1 were used as outgroups. MAG accessions are listed in supplementary table S 5. (B) Host phylogeny based on published mitochondrial cytochrome oxidase subunit I (COI) sequences. *“B.” childressi* was used as an outgroup. Sequence accessions are listed in the supplement (“1.3 Reconstruction of *Bathymodiolus* phylogeny”) (C) Phylogeny of *Bathymodiolus* SOX symbionts from vents on the northern Mid-Atlantic Ridge, based on 171 single-copy marker genes. Colours correspond to vent sites shown in the map. Hybrid individuals from Broken Spur are marked with a grey star. *Bathymodiolus* SOX symbionts from the vent sites Clueless (5°S) and Lilliput (9°S) were used as outgroups. *B.: Bathymodiolus*, MAR: Mid-Atlantic Ridge, GoM: Gulf of Mexico, EPR: East Pacific Rise, J: Japan, NP: North Pacific.

This substantiates previous analyses based on sequencing of the 16S rRNA gene and internal transcribed spacer that these two *Bathymodiolus* species harbour different SOX symbiont subspecies of the same bacterial species (DeChaine et al., 2006; Duperron et al., 2006; Won, Hallam, O’Mullan, Pan, et al., 2003). Our phylogenomic analyses revealed that all *Bathymodiolus* individuals from Broken Spur harboured a third SOX symbiont subspecies (Figure 1 A, C). This new subspecies is most closely related to the *B. puteoserpentis* SOX symbiont subspecies from mussels collected south of Broken Spur. These two symbiont subspecies form a sister group to the SOX symbiont subspecies of *B. azoricus* collected at vents north of Broken Spur.

To evaluate if the SOX symbionts of Broken Spur parental and hybrid *Bathymodiolus* differed, we compared their average nucleotide identities (ANI) and estimated genomic differentiation (F_ST_) based on ∼2500 orthologous genes. We found no significant differences, and also did not see an effect of the year in which the mussels were collected (Figure 2). Our analyses of SNPs per individual gene revealed that not even one of the ∼2500 orthologous genes had significantly differing F_ST_ values (Mann-Whitney U test of F_ST_ per gene between versus within symbionts of hybrids and parental mussels). Similarly, there was also no significant difference between hybrids and parental individuals in the abundance of symbiont genes (based on a general linear model and Kruskal-Wallace test in ALDEx2 using Benjamini-Hochberg corrected p-value < 0.05) or their presence/absence. These results indicate that the composition and gene repertoire of SOX symbionts in Broken Spur mussels is highly similar or identical in hybrids and parental *B. puteoserpentis*. A study of SOX symbionts in hybrids of *B. thermophilus* and *B. antarcticus* at 23°S in the eastern Pacific also found that these could not be distinguished from parental mussels, based on PCR analyses of seven bacterial marker genes in five parental and three hybrid individuals (Ho et al., 2017).

**Figure 2.**
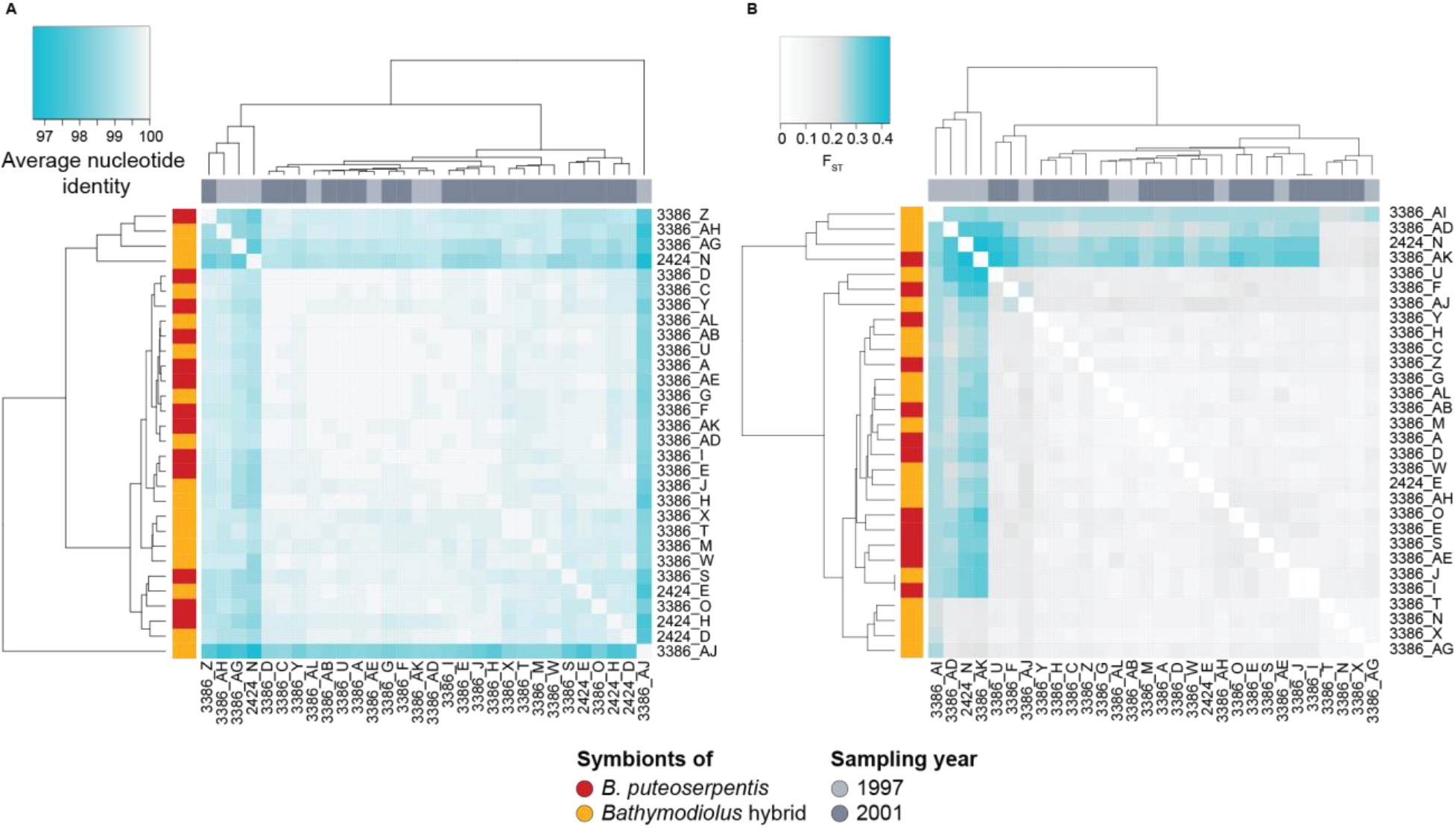
Genome-wide differentiation of *Bathymodiolus* SOX symbionts at Broken Spur based on (A) pairwise average nucleotide identity, and (B) pairwise average F_ST_ based on 2496 orthologous genes. Colour bars represent host genotypes (red: *B. puteoserpentis*, yellow: hybrids) and the sampling year (light grey: 1997, dark grey: 2001). Turquoise indicates a higher differentiation or more dissimilar genomes. Neither clustering based on ANI (A), nor F_ST_ (B) correlates with host genotype (A: r = 0.054, p = 0.222, B: r = 0.006, p = 0.435) or sampling year (A: r = −0.191, p = 0.949, B: r = 0.105, p = 0.150).

Our results raise the question at what level of genetic divergence between two host species differences in their symbiont communities evolve. *B. brooksi* and *B. heckerae*, which regularly co-occur in the Gulf of Mexico, harbour different symbiont species that are only distantly related to each other (Figure 1 A, B). These two mussel species have an estimated splitting time of 15.4 Mya (Faure et al., 2015), and are not known to hybridise. More closely related hosts, such as *B. thermophilus* and *B. antarcticus* (estimated splitting time of 2.5 – 5.3 Mya (Won, Young, Lutz, & Vrijenhoek, 2003)), and *B. azoricus* and *B. puteoserpentis* (estimated splitting time of 8.4 Mya (Faure et al., 2015)), produce fertile hybrids (Johnson et al., 2013; O’Mullan et al., 2001), and have genetically indistinguishable symbionts in zones where they hybridise. This suggests that specificity at the symbiont species level in these horizontally transmitted symbioses evolves only after extended divergence times of tens of millions of years, during which these hosts become genetically dissimilar enough to evolve specific symbiont selection mechanisms.

While *Bathymodiolus* mussels on the northern MAR host the same SOX symbiont species, our phylogenomic analyses revealed clear genetic differentiation in three SOX symbiont subspecies: *B. azoricus, B. puteoserpentis* and Broken Spur subspecies (Figure 1). To better understand the factors that drive this symbiont differentiation, we tested which influence host species, geographic distance, vent type (basaltic versus ultramafic rock) and depth have on symbiont allele frequencies. All variables were highly collinear. For example, the water depth of the vents studied here increases with geographic distance, from 800 m at 37.8°N, to 3050 m at 14.7°N (only the southernmost vent at 13.5°N and 2320 m depth interrupted this pattern). Of the 49 % of the SOX symbiont differentiation that could be explained, host species, depth and vent type explained 23 %, 20 % and 14 % respectively. Geographic distance had by far the strongest effect with 45 % of variation explained (p-value < 0.001, Figure 3).

**Figure 3.**
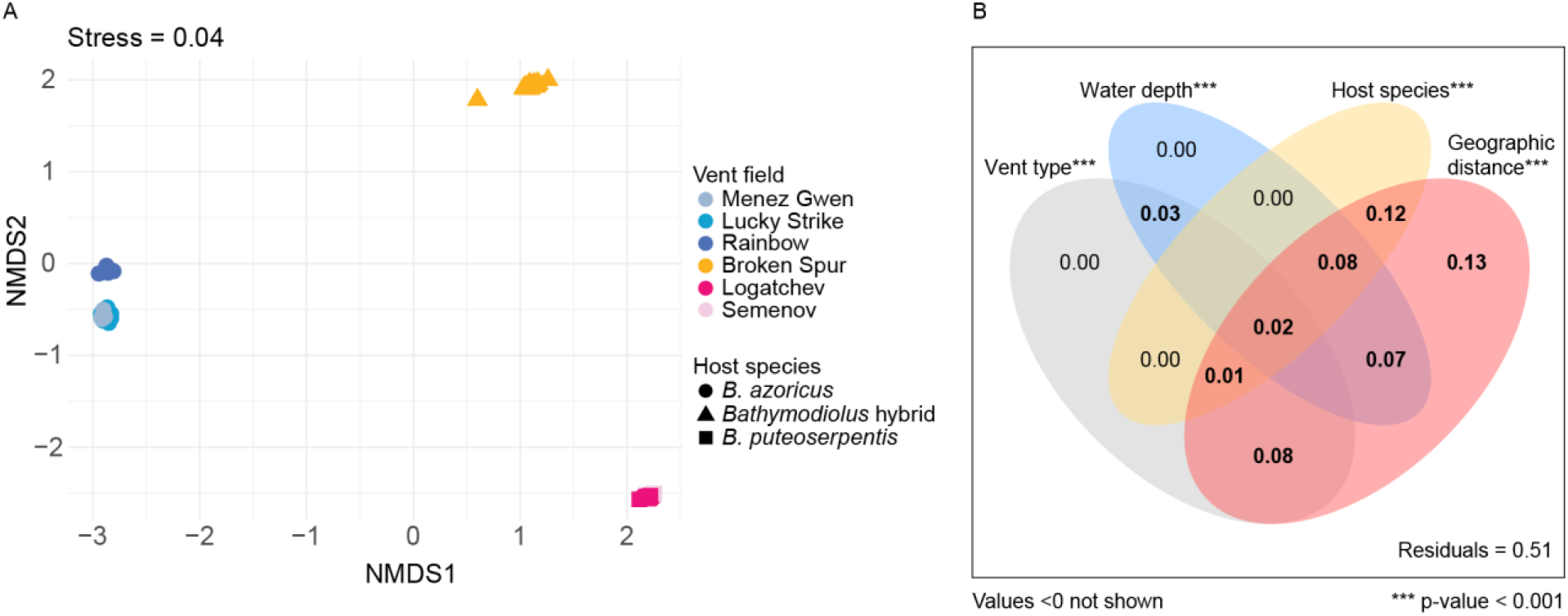
Differentiation of *Bathymodiolus* SOX symbionts at the northern Mid-Atlantic Ridge and the influence of geographic distance, host species and environmental parameters (vent type and water depth). (A) NMDS plot of SOX symbiont allele frequencies show clear separation of the three symbiont subspecies at the northern MAR: *B. azoricus, B. puteoserpentis* and Broken Spur symbiont subspecies. Symbionts from Broken Spur cluster together regardless of the species affiliation of their host (hybrid versus *B. puteoserpentis)*. Colours correspond to vent fields, shapes to host species. (B) Variation partitioning of explanatory variables used in the RDA (Supplementary Figure S 3). The variables vent type, water depth, host species and geographic distance explain 49 % of the total variation, with 45 % of the variation explained by geographic distance. P-values are based on permutation tests with 1000 repetitions.

There are at least two explanations for why geographic distance has such a large effect on the SOX symbiont composition of *Bathymodiolus* mussels from the northern MAR. The first is that genetic differences between the hosts increase with geographic distance. However, population genetic analyses of *B. azoricus* and *B. puteoserpentis* from the same vents as in our study indicated no genetic structuring within each of these host species (Breusing et al., 2016). This indicates that host genetics do not play a major role in structuring the SOX symbiont composition. The second, more likely explanation is that the free-living pool of SOX symbionts is geographically structured. *Bathymodiolus* mussels acquire their symbionts horizontally from the environment, presumably when the larvae settle on the seafloor (Won, Hallam, O’Mullan, Pan, et al., 2003), and must therefore take up the free-living symbionts present in the surrounding water. At Broken Spur, hybrids and *B. puteoserpentis* host genetically indistinguishable symbionts, and these differ from the symbionts of *B. azoricus* and *B. puteoserpentis* from vent sites to the north and south of Broken Spur. This indicates that in these two closely related host species, geographic location but not host genetics drives the composition of their SOX symbiont communities.

Understanding the biogeography of the free-living stages of microbial symbionts and other as yet uncultured microorganisms is currently one of the biggest challenges in microbial ecology. While there is evidence that ‘everything is everywhere, but the environment selects’ (Baas Becking, 1934; Wit & Bouvier, 2006), there is also increasing data showing that dispersal limitation shapes the biogeography of marine microorganisms (Dick, 2019; Martiny et al., 2006). Almost nothing is known about the biogeography of uncultivable marine microorganisms at the subspecies or strain level, as most species are rarely abundant enough to allow phylogenetic analyses at such high resolution. Advances in high-throughput short-read, and particularly long-read sequencing, coupled with bioinformatic methods for revealing genetic structuring of microbial populations, are now providing us with the tools for resolving the intraspecific diversity of environmental microorganisms. Our study highlights the importance of gaining a better understanding of the free-living community of microbial symbionts to disentangle the genetic, environmental and geographic factors that contribute to the ecological and evolutionary success of animal–microbe associations in which the symbionts are acquired from the environment.

## Data availability

Sequence data (metagenomes and symbiont MAGs) are available in the European Nucleotide Archive (ENA) at EMBL-EBI under project accession number PRJEB36976 (https://www.ebi.ac.uk/ena/data/view/PRJEB36976). The data, together with their metadata, were deposited using the data brokerage service of the German Federation for Biological Data (GFBio (Diepenbroek et al., 2014)), with the standard information on sequence data provided as recommended (Yilmaz et al., 2011).

## Acknowledgements

We thank the captains, crews and funding agencies of the sampling cruises AT-03 and AT-05, and the Monterey Bay Aquarium Research Institute and R. C. Vrijenhoek for providing samples. We are grateful to T. Reusch, J. Dierking, K. Trübenbach and P. Weist for the opportunity to perform host genotyping at the GEOMAR and their assistance in the laboratory. We also thank J. Wippler for scientific input, T. Enders for troubleshooting support, and A. Kupczok, L. G. E. Wilkens, and B. Geier for comments on the manuscript. This work was funded by the Max Planck Society, the MARUM German Research Foundation (DFG) Research Centre/Clusters of Excellence “The Ocean in the Earth System” and “The Ocean Floor – Earth’s Uncharted Interface” at the University of Bremen, an European Research Council Advanced Grant (BathyBiome, 340535), a Gordon and Betty Moore Foundation Marine Microbiology Initiative Investigator Award to ND (GBMF3811) and a National Science Foundation Grant (NSF OCE-1736932) to Roxanne Beinart (mentor of CB).

## Contributions

MÜ, RA, LS and ND conceived the study. MÜ performed laboratory work and analyses of symbionts and hosts, prepared figures and tables, submitted data and code, and wrote the initial draft. YS, RA and LS contributed to analyses of the symbionts. CB provided samples and contributed to analyses of the host. MÜ, RA, YS and ND interpreted the results with advice from the other co-authors. MÜ, RA, YS and ND revised the final manuscript with input from all co-authors.

## Supplementary information

Supplementary information is available at https://www.biorxiv.org.

