## Supplementary Information for "Deep-sea mussels from a hybrid zone on the Mid-Atlantic Ridge host genetically indistinguishable symbionts"

#### 1 Supplementary materials & methods

##### 1.1 Sampling, DNA extraction and metagenomic sequencing

Mussels were sampled at Broken Spur (29°10.0'N, 43°10.0'W at 3045 – 3056 m water depth) with the deep-sea submersible Alvin during the Atlantis cruises AT-03/03 (1997) and AT-05/03 (2001). Upon recovery on board, small mussels (< 25 mm) were frozen whole at -70°C, while larger mussels were dissected before freezing at -70°C (Won et al., 2003). An overview map of sites analysed in this study was plotted with RStudio v1.3.959 using R v3.6.3 and the packages rnaturalearth v0.1.0, legendMap v1.0 and ggplot2 v3.3.0 (Andy South, 2017; Ewen Gallic, 2016; RStudio Team, 2015; Wickham et al., 2019).

Genomic DNA was extracted from gill (metagenomic libraries 3386-A~AL) or a combination of gill, mantel and digestive tissue (called mixed tissue hereafter, metagenomic libraries 2424-A~O) depending on sample availability. DNA extractions were performed with either the AllPrep DNA/RNA/Protein MiniKit (Qiagen, Hilden, Germany) or DNAeasy Blood & Tissue kit (Qiagen, Hilden, Germany) according to the manufacturer's protocols with the following modifications: Prior to extraction, frozen sample pieces (5-10 mm) were homogenised by bead beating in MP Biomedicals Lysing Matrix B using an MP Biomedicals FastPrep-24 (Thermo Fisher Scientific, Waltham,

USA) for 30 s at 6.5 m/s. In the elution step, samples were incubated for 10 min at room temperature before centrifugation. Volumes of eluent were halved, and elution was repeated with the first eluate to maximise DNA yields. Metagenomic libraries were generated with the Nextera DNA Flex Library Prep Kit (Illumina, San Diego, CA, USA) and the Illumina TruSeq DNA Samples Prep Kit (BioLABS, Frankfurt, Germany). Library preparation and sequencing of 150 bp paired-end metagenomic reads were performed by the Max Planck-Genome-centre Cologne, Germany (https://mpgc.mpipz.mpg.de/home/) on HiSeq 2500 or 3000 machines. Details of sampling, DNA extraction and sequencing of all samples used in this study are summarised in Supplementary Table S 1.

##### 1.2 Identification of hybrid host individuals

Mussels were genotyped based on 18 species-diagnostic single-nucleotide polymorphism (SNP) markers and identified as hybrid or parental species with subsequent bioinformatic analyses using 1) STRUCTURE v2.3.4 with strauto v1.0 and CLUMPAK (http://clumpak.tau.ac.il/), 2) introgress v1.22 in RStudio, and 3) NEWHYBRIDS v1.1 (Anderson & Thompson, 2002; Chhatre & Emerson, 2017; *CLUMPAK Server*, n.d.; Falush et al., 2003; Gompert & Buerkle, 2009, 2010; Hubisz et al., 2009; Pritchard et al., 2000). The analysis is based on the method developed in (Breusing et al., 2016, 2017). Results of all three programmes are shown in Supplementary Table S 2. Classification based on introgress was used as the basis for analyses of mussel symbionts, as it had low misidentification rates for hybrid and parental species as reported in (Breusing et al., 2017). Since not all programmes supported the classification of backcrosses and their identification was less reliable than for hybrid and parental species in (Breusing et al., 2017), these samples were excluded from further analyses.

##### 1.3 Reconstruction of *Bathymodiolus* phylogeny

Sequences of the mitochondrial marker gene cytochrome c oxidase subunit I or full mitochondrial genomes were downloaded from NCBI (database accessed 2020-02-11) for *B. azoricus* (LN833437), *B. brooksi* (KU597634), *B. heckerae* (KU659139), *B. puteoserpentis* (KU597632), *B.* sp. Lilliput (LN833440), *B.* sp. Clueless (LT674164), *B. septemdierum* (AP014562), *"B." childressi* (ANY30357) and *B. thermophilus* (MK721544) (NCBI Resource Coordinators, 2016). We aligned the sequences with MUSCLE v3.8.31 (Edgar, 2004a, 2004b), and reconstructed a phylogenetic tree using IQTREE v1.6.9 with 1000 samples for ultrafast bootstrap and the mtZoa model, which was selected as the best model by Model Finder based on the Bayesian Information Criterion (Kalyaanamoorthy et al., 2017; Minh et al., 2013; Nguyen et al., 2015; Rota-Stabelli et al., 2009). The tree was visualised with iTol v5.5 (Letunic & Bork, 2019) and edited with Adobe Illustrator 2020 (Adobe, 2020).

##### 1.4 Metagenome assembly and symbiont binning

Metagenomic reads were adapter-trimmed and quality-filtered to a PHRED score of 2 with BBDuk, merged with BBMerge, and error-corrected and normalized to 80x average coverage with BBNorm from BBTools v37.28 (Bushnell, 2014). Merged and unmerged reads were assembled with Megahit v1.0.3 using a maximum k-mer size of 127 (D. Li et al., 2015, 2016). Initial metagenome-assembled genomes (MAGs) were obtained with Metabat2 v2.10.2 automated binning, after mapping with BBMap, and sorting of bam files with samtools v1.9 (Kang et al., 2015; H. Li et al., 2009). MAGs were identified based on small subunit ribosomal RNA gene sequences (SSUs, detected by barrnap v0.6 and classified with vsearch v2.6.2 against the SILVA SSU database v132) and other taxonomic marker genes (detected by Amphora2), through visualisation with gbtools v2.6.0 in RStudio (Edgar, 2010; Quast et al., 2013; Rognes et al., 2016; Seah & Gruber-Vodicka, 2015; Seemann, 2014a; Wu & Scott, 2012). This metagenomic workflow was initially performed separately for gill and mixed tissue. If both sample types were available for the same mussel individual, we confirmed that their symbiont MAGs were identical based on GC content, coverage, and average nucleotide identity (ANI) before pooling the reads and repeating the workflow to increase symbiont coverage. Similarly, if Metabat2 split the sulphur-oxidising (SOX) symbiont sequences into multiple bins, these were pooled after assessment of their GC content, coverage and ANI. Completeness, contamination and strain heterogeneity of MAGs were estimated throughout the binning process with CheckM v1.0.17 based on 280 gammaproteobacterial marker genes (*HMMER*, n.d.; Hyatt et al., 2010; Matsen et al., 2010; Parks et al., 2015). Due to the presence of multiple strains in the symbiont population (Ansorge et al., 2019), we expected duplicates of marker genes with highly similar sequences, such as the ones reported by CheckM as strain heterogeneity. We therefore corrected contamination rates and calculated the contamination that could not be attributed to strain heterogeneity.

For MAGs with a completeness below 90 %, an additional step of assembly and binning was performed to yield MAGs with higher completeness. In such cases, the incomplete bin was used as a reference for mapping to recruit symbiont reads for assembly with SPAdes v3.12.0 (Bankevich et al., 2012). These draft genome assemblies were manually binned in Bandage v0.8.1 based on sequence nodes connected in the assembly graph (Wick et al., 2015).

Additional statistics of symbiont MAGs were calculated using the stats.sh script of BBTools (Supplementary Table S 3). Read coverage of symbiont MAGs was estimated with samtools after mapping raw reads against MAGs with BBMap. High-quality symbiont MAGs with a completeness above 90% and a contamination below 5% (after correction for strain heterogeneity) were used for further analysis with one exception: The MAG of library 3386_H was 89 % complete, only <1 % less than the cutoff, but had no contamination.

#### 1.5 Analyses of SOX symbionts based on symbiont MAGs

All analyses of SOX symbionts conducted in this study, their input data and the level of resolution are summarised in Supplementary Table S 4.

##### *1.5.1 Phylogenomic analysis of SOX symbionts*

To investigate the phylogeny of SOX symbionts from mussels in Broken Spur, we constructed a phylogenomic tree with 171 gammaproteobacterial, single-copy marker genes from the SOX MAGs as well as closely-related symbiotic and free-living bacteria. Accession numbers and publications for all reference MAGs/genomes used in the analysis are listed in Supplementary Table S 5.

A protein alignment of the 171 marker genes (Data file: Phylogenomics_NMAR.fasta, available at https://github.com/muecker/Symbionts_in_a_mussel_hybrid_zone) was obtained with GToTree v1.4.11 (Capella-Gutiérrez et al., 2009; Edgar, 2004a, 2004b; *HMMER*, n.d.; Hyatt et al., 2010; Lee, 2019). The alignment was visually inspected in Geneious v11.1.5 (Kearse et al., 2012). One gene sequence (of sample 1586K) had a high proportion of mismachtes to all other sequences and was identified as contamination by blasting against the NCBI database. This sequence was removed from the alignment. Using the LG+F+R6 amino acid model ((Le & Gascuel, 2008), best model according to Model Finder), and 1000 samples for ultrafast bootstrap, we reconstructed a phylogenomic tree of SOX symbionts and their closest relatives, and edited it with iTol and Adobe Illustrator.

Correlation between symbiont $F_{ST}$ and geographic distance was tested using the Mantel test. The multiple sequence alignment was imported into R with the read.alignment function of R package seqinr v3.4-5 and transformed into a genind object using the alignment2genind function of adegenet v2.1.1 (Charif et al., 2019; Jombart et al., 2020). We caluclated the pairwise $F_{ST}$ of symbionts between all locations on the northern MAR using pairwise.fst of R package hierfstat v0.04-33 (Goudet & Jombart, 2015). Geographic distances between vent sites were calculated based on the coordinates using https://www.movable-type.co.uk/scripts/latlong.html, and transformed into a dist object using the R function dist. To account for the geographic subdivision of the host species that can also cause patterns similar to isolation-by-distance (Meirmans, 2012), we performed a stratified Mantel test using the mantel function of R package vegan v2.5-5, and the host species groups as strata (mantel(FST, geo_distances,strata=groups_NMAR) (Oksanen et al., 2019). After a statistical significant result, the $F_{ST}$ values between symbionts from different sites were plotted against their geographic distances for visual.

##### *1.5.2 Average nucleotide identity of SOX symbionts*

To analyse how similar symbiont MAGs from Broken Spur were to each other, we analysed their pairwise ANI values of the aligned fraction (0.48 – 0.99%) with fastANI v1.1 (Jain et al., 2018) (Data file: Average_nucleotide_identity_SOX_symbionts_Broken_Spur.csv). Samples were clustered and represented based on their average ANI values in a heatmap with dendrogram, generated in RStudio using the packages gplots v3.0.1.1 and maditr v0.6.2 (Demin, 2019; Warnes et al., 2019).

Correlation of SOX symbiont ANI values and sampling year was tested with a Mantel test (mantel function of R package vegan, 5039 permutations) using the Spearman's rank correlation coefficient after transforming sampling year information into euclidean distances. To test the correlation between SOX symbiont ANI values and host genetics, pairwise genetic distances between host individuals were calculated based on 18 species-diagnostic SNP markers (see "1.2. Identification of hybrid host individuals"). Host SNP markers were imported to RStudio and converted into a dataframe using the read.structure and genind2df functions of the package adegenet. We subsequently calculated pairwise genetic distances between individuals with the dist.gene function of R package ape v5.3 (Paradis et al., 2019). Correlation was tested as described above for ANI values vs. sampling year.

#### 1.6 Analyses of symbiont population based on single-nucleotide polymorphisms

Recent advances in whole-(meta)genome approaches have increased resolution and sensitivity of analyses, and advanced our knowledge on strain diversity in deep-sea mussel symbiont populations (Ansorge et al., 2019; Ikuta et al., 2016; Picazo et al., 2019). We therefore performed genome-wide SNP analyses of SOX symbionts based on 2496 orthologous genes to investigate symbiont population differentiation between different mussel individuals from Broken Spur.

We used a gene catalogue of 3204 orthologues (see below) as a reference for SNP identification. The catalogue was annotated using prokka v1.1, resulting in 2496 genes with annotation (including hypothetical proteins) (Seemann, 2014b). Genes without any annotation, mostly short (<300 bp), probably fragmented genes, were excluded from the analysis. SNP calling was performed as described in (Ansorge et al., 2019) using scripts available at https://github.com/rbcan/MARsym_paper with a few adjustments to newer software versions. In summary, raw reads were adapter trimmed and quality filtered to a PHRED score of 20 and subsequently mapped to the reference with a minimum identitiy of 95 % using BBMap. We realigned reads around indels and downsampled to an average coverage of 70x. Samples that did not meet the coverage threshold were excluded from the analysis. The steps above were performed with samtools, Picard tools v1.1.02 and the Genome Analysis Toolkit (GATK) v3.7-0 (Broad Institute, 2013; H. Li et al., 2009; McKenna et al., 2010). SNPs were called with GATK HaplotypeCaller, and unreliable SNPs were filtered with GATK VariantFiltration (settings: QD < 2; FS > 60; MQ < 40, MQRankSum < -20, ReadPosRankSum < -8). Spearman's rank correlation coefficient was calculated in RStudio using the R function cor to test for correlation of SNP density (#SNPs/kb) with shell size. The fixation index $F_{ST}$ was calculated for each gene with a script (https://github.com/deropi/BathyBrooksiSymbionts/tree/master/Population_structure_analyses) previously used in (Picazo et al., 2019), and averaged per host individual (Data file: Pairwise_mean_FST_SOX_symbionts_Broken_Spur.csv). Mean pairwise $F_{ST}$ values were plotted in a heatmap, and correlation between $F_{ST}$ and sampling year and $F_{ST}$ and host genotype was tested as described above for ANI values.

#### 1.7 Analysis of differences in gene repertoire between symbionts from hybrids and parental species

##### *1.7.1 Gene presence/absence and abundance analyses*

To examine whether there are differences in the gene repertoire of the symbiont populations between hybrid and parental mussels, we analysed the presence/absence of genes specific to either group of mussels and their relative abundances. We annotated all

MAGs with prokka and clustered orthologues with GET_HOMOLOGUES v3.2.3 using the OrthoMCL algorithm (Altschul et al., 1997; Brown et al., 1998; Buchfink et al., 2015; Contreras-Moreira & Vinuesa, 2013; Finn et al., 2016; *HMMER*, n.d.; Kristensen et al., 2010; L. Li et al., 2003; Stajich et al., 2002; Vinuesa & Contreras-Moreira, 2015), resulting in a gene catalogue of 3204 orthologues (Data file: Orthologue_gene_catalogue_OMCL_SOX_symbionts_Broken_Spur.fasta; also used in SNP-identification above). Using the parse_pangenome_matrix.pl script of GET_HOMOLOGUES, we tested for genes that were present in at least 90 % of symbiont genomes from *B. puteoserpentis* and absent in at least 90 % of symbiont genomes from hybrids, and vice versa.

To further analyse gene abundances, raw reads of all libraries were mapped to the orthologous gene catalogue with BBMap and downsampled to 70x coverage with samtools. The fasta sequences were extracted from downsampled bam files using samtools and pseudoaligned to the catalogue using kallisto v0.46.0 (Bray et al., 2016; Bushnell, 2014). The gene coverage was estimated using the abundance_estimates_to_matrix.pl script of Trinity v2.5.1 (Grabherr et al., 2011; Haas et al., 2013) (Data file: Gene_counts_kallisto_SOX_symbionts_Broken_Spur.matrix).

To account for the compositionality of the data, the gene abundances were statistically evaluated using ALDEx2 v1.16.0 and data.table v1.12.2 in RStudio (Dowle et al., 2019; Fernandes et al., 2013, 2014; Gloor et al., 2016; RStudio Team, 2015). We used the aldex.clr module to prepare the data using host categories (hybrid or *B. puteoserpentis*) as condition. With the aldex.kw command, we ran a general linear model and a Kruskal Wallace test for one way ANOVA.

##### *1.7.2 Analysis of gene differentiation between symbionts from hybrids and parentals*

We analysed population differentiation ($F_{ST}$) based on SNP frequencies in 2496 orthologue genes to find genes that have higher differentiation ‘between’ symbionts of hybrids and parental species than variations ‘within’ symbionts of the same host category (hybrids or *B. puteoserpentis*, Supplementary Figure S 1). $F_{ST}$ values were acquired as described above (1.6 “Analyses of symbiont population based on single-nucleotide polymorphisms”) and reformatted for ‘between’ versus ‘within’ statistical comparisons (Data file: Per_gene_FST_SOX_symbionts_Broken_Spur.zip). We used a Mann–Whitney U test in RStudio to test the null hypothesis that there is no significant difference between $F_{ST}$ of genes in the ‘between’ (hybrids versus parental species) and the ‘within’ (hybrids versus hybrids, parental versus parental species) categories. To reduce false discovery rates, the test was repeated with a dataset of random $F_{ST}$ values. More genes with p-value <0.05 were detected in the random dataset than in the actual data, and no p-values of the real dataset were below those from the random dataset.

This indicates that all genes detected as significant (p<0.05) for the real data can be attributed to type I error, and that there was no gene more differentiated between symbionts of hybrid and parental mussels than within the same host category.

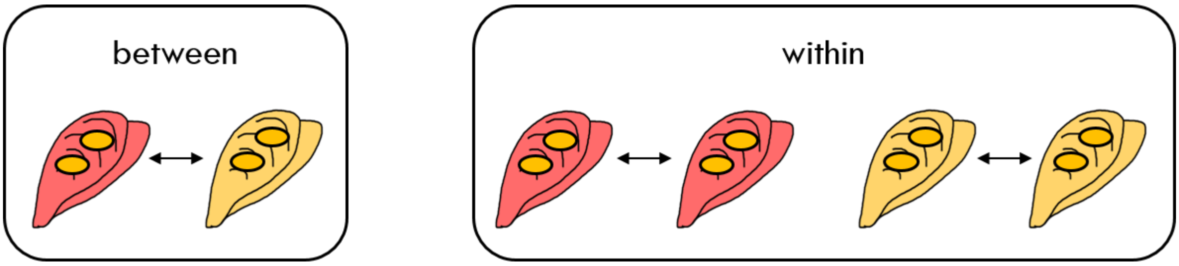

**Supplementary Figure S 1 | Categories for per gene $F_{ST}$ analysis.** Between: Comparison of SOX symbionts from *B. puteoserpentis* (red) and hybrid mussels (yellow). Within: Comparison among SOX symbionts from *B. puteoserpentis* and among SOX symbionts from hybrid mussels.

#### 1.8 Redundancy analysis of SOX symbiont allele frequencies from *Bathymodiolus* mussels along the northern Mid-Atlantic Ridge

Redundancy analysis (RDA) allows to eliminate redundant information in genetic data and its associations with environmental variables, and to assess the proportion of variation explained by these environmental variables (Borcard et al., 2018). We performed an RDA in RStudio using the vegan package to test how much of the variation in symbiont allele frequencies can be explained by geographic distance, the vent type (basaltic versus ultramafic rock), the associated host species and depth. In hydrothermal systems, the composition of host rock has been described to influence the chemical composition and thereby the energy available for chemoautotrophs in vent fluids (Amend et al., 2011) which is why we chose vent type as environmental parameter. Depth and vent type were retrieved from the InterRidge Vents Database v3.4 (https://vents-data.interridge.org/, accessed 2020-06-15). As a reference for SNP analysis, we constructed a gene catalogue based on all SOX symbiont MAGs from the northern MAR (Data file: Orthologue_gene_catalogue_OMCL_SOX_symbionts_NMAR.fasta) using the workflow described above (1.7.1 “Gene presence/absence and abundance analyses”). To obtain allele frequencies, we performed a SNP analysis based on the gene catalogue of all SOX symbionts from the northern MAR as described above (1.6 “Analyses of symbiont population based on single-nucleotide polymorphisms”). We extracted the AD (read depth per allele) and DP (read depth) field from VCF files using GATK’s VariantToTable tool, divided AD by DP to obtain allele frequencies per sample and merged the individual tables using join on the Linux command line (Data file: Allele_frequencies_SOX_symbionts_NMAR.csv). For the RDA, site coordinates were scaled and computed as orthogonal polynomials with R package stats v3.6.3 (function poly) as suggested by (Borcard et al., 2018; Legendre, 1993; Ter Braak, 1987). We performed a forward selection on the polynomials with the ordistep function of R package vegan, ran the RDA with all variables and calculated an adjusted $R^2$. We assessed the significance of the RDA, the individual axes and the explanatory variables with the vegan package function anova.cca using 1000 permutations. To explore how much variation can be explained by each explanatory variable, we performed a variation partitioning using the varpart function of vegan and plotted it with R base function plot. The RDA triplot was plotted using the ggord package v1.1.4 (Marcus W. Beck, 2019) (Supplementary Figure S 3). As overview visualisation of the allele frequencies, we calculated an NMDS using the metaMDS function of vegan and plotted it with ggplot2. All figures were modified with Adobe Illustrator.

### 2 Supplementary results & discussion

#### 2.1 *B. puteoserpentis* and hybrid individuals identified in Broken Spur

We genotyped mussels from Broken Spur and identified *B. puteoserpentis* and hybrid individuals. *B. azoricus* mussels were not detected in all methods used, except for one mussel (3676-15/3386_N) that was identified as *B. azoricus* by NEWHYBRIDS. However, this result was not supported by the two other programmes, suggesting that the mussel is more likely a hybrid.

The absence of *B. azoricus* in Broken Spur could be due to bathymetric limitation, as *B. azoricus* usually occurs at shallower depths. Another possible explanation is that the actual hybrid zone might be further north as suggested by (Breusing et al., 2017). Lastly, it cannot be ruled out that *B. azoricus* mussels were not found during sampling as the number of mussels collected was limited and the distribution rather scattered at Broken Spur.

All hybrids identified by INTROGRESS were in the F2 to F4 generation indicating that the hybrids are fertile. Although the exact status of backcrosses, especially which generations of backcrosses are actually present, was uncertain, multiple mussels were identified as backcrosses by NEWHYBRIDS and INTROGRESS. Together with the admixture values reported by STRUCTURE, this suggests that there is still gene flow between hybrids and the populations of parental species.

##### 2.2 No difference in gene abundances between symbionts from hybrids and parental species

We analysed 3204 orthologous genes to detect genes that are exclusive to either symbionts of hybrid or symbionts of *B. puteoserpentis*. GET_HOMOLOGUES detected none of such genes, even with lower stringency (presence in >90 % in one and <90 % in the other group).

When comparing gene abundances between symbionts from hybrid and *B. puteoserpentis* mussels using the statistical analysis with ALDEx2, no genes were significantly different in their abundances (Benjamini-Hochberg corrected p-value < 0.05). Functional variation has previously been shown to occur among symbiont populations from different vents along the MAR (Ansorge et al., 2019). However, the gene repertoire of SOX symbiont populations within Broken Spur did not vary according to host genotype, indicating that hybrids and parental species do not select their symbionts based on different functions.

##### 2.3 No correlation of SNPs/kb with mussel shell size

Picazo et al. (2019) previously detected lower strain diversity in large (146–241 mm) compared to medium-sized (72–141 mm) mussels in the Gulf of Mexico that might be explained by self-infection and slower symbiont uptake in older mussels (Picazo et al., 2019). We did not detect any correlation of SNPs/kb with shell size, which might be due to the relatively limited size range of the analysed mussels (24–133 mm).

##### 2.4 Isolation-by-distance of *Bathymodiolus* SOX symbiont subspecies at the northern MAR

Phylogenomic analysis of 171 gammaproteobacterial marker genes revealed that symbiont genetic variation ($F_{ST}$ based on the amino acid alignment) was positively correlated with geographic distance (r = 0.7471, p = 0.035). However, a gradual genetic change along a geographic gradient as would be expected under an evolutionary isolation-by-distance (IBD) model (Wright, 1943) could not be observed (Supplementary Figure S 2). Mantel tests are often used to test for isolation-by-distance which is why we included this analysis here. However, its use has been discouraged (Meirmans, 2015). We therefore used a redundancy analysis in this study (Figure 3, Supplementary Figure S 3, Supplement 1.8 “Redundancy analysis of SOX symbiont allele frequencies from *Bathymodiolus* mussels along the northern Mid-Atlantic Ridge” and main text).

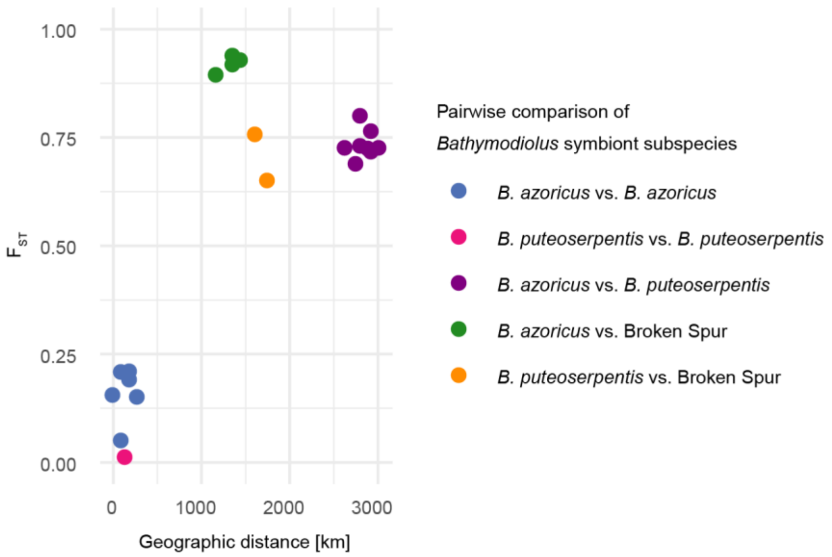

**Supplementary Figure S 2 | Relation of $F_{ST}$ and geographic distance between *Bathymodiolus* SOX symbiont populations from different vent fields along the northern MAR.** Displayed $F_{ST}$ values are averaged pairwise $F_{ST}$ between symbionts from all mussels at a vent field. Each dot represents one pairwise comparison between two sites, comparisons of a site with itself are not shown. Colours correspond to the comparisons of the different symbiont subspecies (*B. azoricus* type, present at Menez Gwen – White Flames, Lucky Strike – Montsegur, Lucky Strike – Eiffel Tower and Rainbow; *B. puteoserpentis* type, present at Logatchev Quest and Semenov; Broken Spur type, present at Broken Spur).

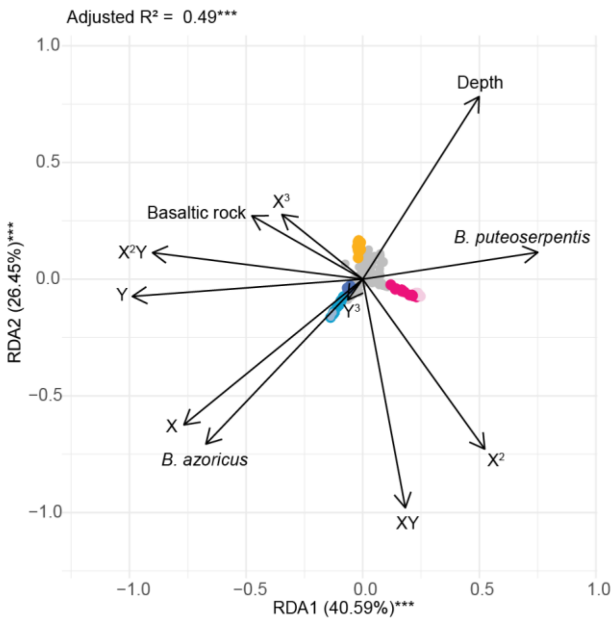

**Supplementary Figure S 3 | Influence of geographic distance, host species and environmental parameters on differentiation of *Bathymodiolus* SOX symbionts at the northern MAR.** Redundancy analysis triplot (scaling 2, wa scores) showing the influence of geographic distance (forward selected variables X, Y, $X^2$, $X^3$, $X^2Y$, $Y^3$ represent orthogonal polynomials of latitude and longitude), host species (*B. azoricus* and *B. puteoserpentis*), the vent type (only basaltic rock displayed) and water depth on symbiont allele frequencies. *** p-value < 0.001. P-values are based on permutation tests with 1000 repetitions.

**Supplementary Table S 1 | Sample overview.** ID consists of dive number and mussel ID. Shell length (size) is listed when data were available according to cruise material in http://dlacruisedata.whoi.edu/AT/AT003L03/ (accessed 2019-05-29). AllPrep: AllPrep DNA/RNA/Protein MiniKit (Qiagen); DNAeasy: DNAeasy Blood & Tissue kit (Qiagen), Nex: Nextera DNA Flex Library Prep Kit (Illumina); TruSeq: Illumina TruSeq DNA Samples Prep Kit (BioLABS).

| Cruise | ID | Latitude | Longitude | Depth [m] | Sampling date | Material | DNA extraction | Libprep | Sequencing | Library | Size [mm] |
|---|---|---|---|---|---|---|---|---|---|---|---|
| AT_05/03 | 3676-1 | 29.1672 | -43.1742 | 3045 | July 19, 2001 | gill | AllPrep | Nex | HiSeq 3000 | 3386_A | 119.8 |
| AT_05/03 | 3676-2 | 29.1672 | -43.1742 | 3045 | July 19, 2001 | gill | AllPrep | Nex | HiSeq 3000 | 3386_B | 113.8 |
| AT_05/03 | 3676-3 | 29.1672 | -43.1742 | 3045 | July 19, 2001 | gill | AllPrep | Nex | HiSeq 3000 | 3386_C | 112.2 |
| AT_05/03 | 3676-4 | 29.1672 | -43.1742 | 3045 | July 19, 2001 | gill | AllPrep | Nex | HiSeq 3000 | 3386_D | 116.3 |
| AT_05/03 | 3676-5 | 29.1672 | -43.1742 | 3045 | July 19, 2001 | gill | AllPrep | Nex | HiSeq 3000 | 3386_E | 119.1 |
| AT_05/03 | 3676-6 | 29.1672 | -43.1742 | 3045 | July 19, 2001 | gill | AllPrep | Nex | HiSeq 3000 | 3386_F | 114.6 |
| AT_05/03 | 3676-7 | 29.1672 | -43.1742 | 3045 | July 19, 2001 | gill | AllPrep | Nex | HiSeq 3000 | 3386_K | 107.6 |
| AT_05/03 | 3676-8 | 29.1672 | -43.1742 | 3045 | July 19, 2001 | gill | AllPrep | Nex | HiSeq 3000 | 3386_G | 103.6 |
| AT_05/03 | 3676-9 | 29.1672 | -43.1742 | 3045 | July 19, 2001 | gill | AllPrep | Nex | HiSeq 3000 | 3386_H | 75 |
| AT_05/03 | 3676-10 | 29.1672 | -43.1742 | 3045 | July 19, 2001 | gill | AllPrep | Nex | HiSeq 3000 | 3386_I | 122.9 |
| AT_05/03 | 3676-11 | 29.1672 | -43.1742 | 3045 | July 19, 2001 | mixed & gill | DNAeasy & AllPrep | TruSeq, Nex | HiSeq 2500 & 3000 | 3386_J, 2424_A | 101.9 |
| AT_05/03 | 3676-12 | 29.1672 | -43.1742 | 3045 | July 19, 2001 | gill | AllPrep | Nex | HiSeq 3000 | 3386_L | 114.8 |
| AT_05/03 | 3676-14 | 29.1672 | -43.1742 | 3045 | July 19, 2001 | mixed & gill | DNAeasy & AllPrep | TruSeq & Nex | HiSeq 2500 & 3000 | 3386_M, 2424_B | 81.4 |
| AT_05/03 | 3676-15 | 29.1672 | -43.1742 | 3045 | July 19, 2001 | mixed & gill | DNAeasy & AllPrep | TruSeq & Nex | HiSeq 2500 & 3000 | 3386_N, 2424_C | 92.9 |
| AT_05/03 | 3676-16 | 29.1672 | -43.1742 | 3045 | July 19, 2001 | mixed & gill | DNAeasy & AllPrep | TruSeq & Nex | HiSeq 2500 & 3000 | 3386_O, 2424_F | 12.2 |
| AT_05/03 | 3676-17 | 29.1672 | -43.1742 | 3045 | July 19, 2001 | gill | AllPrep | Nex | HiSeq 3000 | 3386_P | 110.2 |

| | | | | | | | | | | | |
|---|---|---|---|---|---|---|---|---|---|---|---|
| AT_05/03 | 3676-18 | 29.1672 | -43.1742 | 3045 | July 19, 2001 | gill | AllPrep | Nex | HiSeq 3000 | 3386_Q | 89.9 |
| AT_05/03 | 3676-19 | 29.1672 | -43.1742 | 3045 | July 19, 2001 | gill | AllPrep | Nex | HiSeq 3000 | 3386_R | 64.7 |
| AT_05/03 | 3676-20 | 29.1672 | -43.1742 | 3045 | July 19, 2001 | mixed & gill | DNAeasy & AllPrep | TruSeq & Nex | HiSeq 2500 & 3000 | 3386_S, 2424_G | 93.6 |
| AT_05/03 | 3676-21 | 29.1672 | -43.1742 | 3045 | July 19, 2001 | gill | AllPrep | Nex | HiSeq 3000 | 3386_T | 93.2 |
| AT_05/03 | 3676-22 | 29.1672 | -43.1742 | 3045 | July 19, 2001 | gill | AllPrep | Nex | HiSeq 3000 | 3386_U | 81.5 |
| AT_05/03 | 3676-26 | 29.1672 | -43.1742 | 3045 | July 19, 2001 | mixed | DNAeasy | TruSeq | HiSeq 2500 | 2424_D | 31.7 |
| AT_05/03 | 3676-27 | 29.1672 | -43.1742 | 3045 | July 19, 2001 | gill | AllPrep | Nex | HiSeq 3000 | 3386_V | 65.4 |
| AT_05/03 | 3676-28 | 29.1672 | -43.1742 | 3045 | July 19, 2001 | mixed | DNAeasy | TruSeq | HiSeq 2500 | 2424_E | 66.7 |
| AT_05/03 | 3676-29 | 29.1672 | -43.1742 | 3045 | July 19, 2001 | mixed | DNAeasy | TruSeq | HiSeq 2500 | 2424_H | 51.8 |
| AT_05/03 | 3676-30 | 29.1672 | -43.1742 | 3045 | July 19, 2001 | gill | AllPrep | Nex | HiSeq 3000 | 3386_W | 94.4 |
| AT_05/03 | 3676-31 | 29.1672 | -43.1742 | 3045 | July 19, 2001 | gill | AllPrep | Nex | HiSeq 3000 | 3386_X | 64.5 |
| AT_05/03 | 3676-32 | 29.1672 | -43.1742 | 3045 | July 19, 2001 | mixed & gill | DNAeasy & AllPrep | TruSeq & Nex | HiSeq 2500 & 3000 | 3386_Y, 2424_I | 88.8 |
| AT_05/03 | 3676-33 | 29.1672 | -43.1742 | 3045 | July 19, 2001 | mixed & gill | DNAeasy & AllPrep | TruSeq & Nex | HiSeq 2500 & 3000 | 3386_Z, 2424_J | 100.5 |
| AT_05/03 | 3676-37 | 29.1672 | -43.1742 | 3045 | July 19, 2001 | gill | AllPrep | Nex | HiSeq 3000 | 3386_AA | 111.4 |
| AT_05/03 | 3676-38 | 29.1672 | -43.1742 | 3045 | July 19, 2001 | gill | AllPrep | Nex | HiSeq 3000 | 3386_AB | 104.7 |
| AT_03/03 | 3125-1 | 29.1667 | -43.1733 | 3056 | July 17, 1997 | gill | AllPrep | Nex | HiSeq 3000 | 3386_AC | |
| AT_03/03 | 3125-2 | 29.1667 | -43.1733 | 3056 | July 17, 1997 | gill | AllPrep | Nex | HiSeq 3000 | 3386_AD | |
| AT_03/03 | 3125-3 | 29.1667 | -43.1733 | 3056 | July 17, 1997 | gill | AllPrep | Nex | HiSeq 3000 | 3386_AE | |
| AT_03/03 | 3125-4 | 29.1667 | -43.1733 | 3056 | July 17, 1997 | gill | AllPrep | Nex | HiSeq 3000 | 3386_AF | |
| AT_03/03 | 3125-5 | 29.1667 | -43.1733 | 3056 | July 17, 1997 | mixed & gill | DNAeasy & AllPrep | TruSeq & Nex | HiSeq 2500 & 3000 | 3386_AG, 2424_K | |
| AT_03/03 | 3125-6 | 29.1667 | -43.1733 | 3056 | July 17, 1997 | mixed & gill | DNAeasy & AllPrep | TruSeq & Nex | HiSeq 2500 & 3000 | 3386_AH, 2424_L | |

| | | | | | | | | | | |
|---|---|---|---|---|---|---|---|---|---|---|
| AT_03/03 | 3125-7 | 29.1667 | -43.1733 | 3056 | July 17, 1997 | mixed & gill | DNAeasy & AllPrep | TruSeq & Nex | HiSeq 2500 & 3000 | 3386_AI, 2424_M |
| AT_03/03 | 3125-8 | 29.1667 | -43.1733 | 3056 | July 17, 1997 | mixed | DNAeasy | TruSeq | HiSeq 2500 | 2424_N |
| AT_03/03 | 3125-9 | 29.1667 | -43.1733 | 3056 | July 17, 1997 | mixed & gill | DNAeasy & AllPrep | TruSeq & Nex | HiSeq 2500 & 3000 | 3386_AJ, 2424_O |
| AT_03/03 | 3125-10 | 29.1667 | -43.1733 | 3056 | July 17, 1997 | gill | AllPrep | Nex | HiSeq 3000 | 3386_AK |
| AT_03/03 | 3125-11 | 29.1667 | -43.1733 | 3056 | July 17, 1997 | gill | AllPrep | Nex | HiSeq 3000 | 3386_AL |

**Supplementary Table S 2 | Genotyping results of 42 mussel individuals from Broken Spur based on analyses with NEWHYBRIDS, INTROGRESS and STRUCTURE.** ID: Dive number–mussel ID; Lib: Metagenomic library name; Bazo: *B. azoricus*; BC azo: Backcross between *B. azoricus* and hybrid; FX: Hybrid in generation X; BC put: Backcross between *B. puteoserpentis* and hybrid; Bput: *B. puteoserpentis*; Genotype: NEWHYBRIDS' genotype category with highest probability.

| **ID** | **Lib** | **NEWHYBRIDS** | | | | | | | | | | | | | **INTROGRESS** | **STRUCTURE** | |
|---|---|---|---|---|---|---|---|---|---|---|---|---|---|---|---|---|---|
| | | **Bazo** | **BC1 azo** | **BC2 azo** | **BC3 azo** | **BC4 azo** | **F1** | **F2-4** | **BC1 put** | **BC2 put** | **BC3 put** | **BC4 put** | ***Bput*** | **Genotype** | | **Bazo** | **Bput** |
| 3676-1 | 3386_A | 0.00 | 0.00 | 0.00 | 0.00 | 0.00 | 0.00 | 0.00 | 0.00 | 0.00 | 0.07 | 0.09 | 0.84 | Bput | ***B. puteoserpentis*** | 0.00 | 1.00 |
| 3676-2 | 3386_B | 0.00 | 0.00 | 0.00 | 0.00 | 0.00 | 0.00 | 0.23 | 0.13 | 0.36 | 0.27 | 0.01 | 0.00 | BC put | **BC *puteoserpentis*** | 0.19 | 0.81 |
| 3676-3 | 3386_C | 0.00 | 0.00 | 0.00 | 0.00 | 0.00 | 0.00 | 1.00 | 0.00 | 0.00 | 0.00 | 0.00 | 0.00 | F2-4 | **F2-4** | 0.32 | 0.68 |
| 3676-4 | 3386_D | 0.00 | 0.00 | 0.00 | 0.00 | 0.00 | 0.00 | 0.00 | 0.00 | 0.00 | 0.04 | 0.07 | 0.89 | Bput | ***B. puteoserpentis*** | 0.00 | 1.00 |
| 3676-5 | 3386_E | 0.00 | 0.00 | 0.00 | 0.00 | 0.00 | 0.00 | 0.00 | 0.00 | 0.00 | 0.07 | 0.08 | 0.84 | Bput | ***B. puteoserpentis*** | 0.00 | 1.00 |
| 3676-6 | 3386_F | 0.00 | 0.00 | 0.00 | 0.00 | 0.00 | 0.00 | 0.00 | 0.00 | 0.02 | 0.28 | 0.16 | 0.54 | BC put | ***B. puteoserpentis*** | 0.01 | 0.99 |
| 3676-7 | 3386_K | 0.00 | 0.00 | 0.00 | 0.00 | 0.00 | 0.00 | 0.00 | 0.00 | 0.13 | 0.50 | 0.17 | 0.20 | BC put | **BC *puteoserpentis*** | 0.04 | 0.96 |
| 3676-8 | 3386_G | 0.00 | 0.01 | 0.00 | 0.00 | 0.00 | 0.60 | 0.36 | 0.04 | 0.00 | 0.00 | 0.00 | 0.00 | F1 | **F2-4** | 0.37 | 0.63 |
| 3676-9 | 3386_H | 0.00 | 0.00 | 0.00 | 0.00 | 0.00 | 0.00 | 1.00 | 0.00 | 0.00 | 0.00 | 0.00 | 0.00 | F2-4 | **F2-4** | 0.28 | 0.72 |
| 3676-10 | 3386_I | 0.00 | 0.00 | 0.00 | 0.00 | 0.00 | 0.00 | 0.00 | 0.00 | 0.00 | 0.07 | 0.09 | 0.84 | Bput | ***B. puteoserpentis*** | 0.00 | 1.00 |
| 3676-11 | 3386_J | 0.00 | 0.00 | 0.00 | 0.00 | 0.00 | 0.00 | 1.00 | 0.00 | 0.00 | 0.00 | 0.00 | 0.00 | F2-4 | **F2-4** | 0.37 | 0.63 |
| 3676-12 | 3386_L | 0.00 | 0.00 | 0.00 | 0.00 | 0.00 | 0.00 | 0.01 | 0.10 | 0.41 | 0.45 | 0.03 | 0.00 | BC put | **BC *puteoserpentis*** | 0.13 | 0.87 |
| 3676-14 | 3386_M | 0.03 | 0.63 | 0.00 | 0.00 | 0.00 | 0.00 | 0.33 | 0.00 | 0.00 | 0.00 | 0.00 | 0.00 | BC azo | **F2-4** | 0.57 | 0.43 |
| 3676-15 | 3386_N | 0.86 | 0.12 | 0.01 | 0.00 | 0.00 | 0.00 | 0.00 | 0.00 | 0.00 | 0.00 | 0.00 | 0.00 | Bazo | **F2-4** | 0.69 | 0.31 |
| 3676-16 | 3386_O | 0.00 | 0.00 | 0.00 | 0.00 | 0.00 | 0.00 | 0.00 | 0.00 | 0.00 | 0.10 | 0.10 | 0.79 | Bput | ***B. puteoserpentis*** | 0.00 | 1.00 |
| 3676-17 | 3386_P | 0.00 | 0.00 | 0.00 | 0.00 | 0.00 | 0.00 | 0.00 | 0.06 | 0.40 | 0.50 | 0.04 | 0.00 | BC put | **BC *puteoserpentis*** | 0.13 | 0.87 |
| 3676-18 | 3386_Q | 0.00 | 0.00 | 0.00 | 0.00 | 0.00 | 0.00 | 0.00 | 0.00 | 0.04 | 0.28 | 0.14 | 0.53 | Bput | **BC *puteoserpentis*** | 0.01 | 0.99 |
| 3676-19 | 3386_R | 0.00 | 0.00 | 0.00 | 0.00 | 0.00 | 0.00 | 0.00 | 0.05 | 0.38 | 0.52 | 0.06 | 0.00 | BC put | **BC *puteoserpentis*** | 0.12 | 0.88 |
| 3676-20 | 3386_S | 0.00 | 0.00 | 0.00 | 0.00 | 0.00 | 0.00 | 0.00 | 0.00 | 0.01 | 0.21 | 0.15 | 0.63 | Bput | ***B. puteoserpentis*** | 0.00 | 1.00 |
| 3676-21 | 3386_T | 0.00 | 0.02 | 0.00 | 0.00 | 0.00 | 0.31 | 0.66 | 0.01 | 0.00 | 0.00 | 0.00 | 0.00 | F2-4 | **F2-4** | 0.40 | 0.60 |
| 3676-22 | 3386_U | 0.00 | 0.00 | 0.00 | 0.00 | 0.00 | 0.20 | 0.59 | 0.19 | 0.03 | 0.00 | 0.00 | 0.00 | F2-4 | **F2-4** | 0.29 | 0.71 |
| 3676-26 | 2339_D | 0.00 | 0.07 | 0.00 | 0.00 | 0.00 | 0.24 | 0.69 | 0.00 | 0.00 | 0.00 | 0.00 | 0.00 | F2-4 | **F2-4** | 0.43 | 0.57 |

| | | | | | | | | | | | | | | | | | |
|---|---|---|---|---|---|---|---|---|---|---|---|---|---|---|---|---|---|
| 3676-27 | 3386_V | 0.00 | 0.00 | 0.00 | 0.00 | 0.00 | 0.00 | 0.00 | 0.00 | 0.12 | 0.50 | 0.18 | 0.19 | BC put | **BC *puteoserpentis*** | 0.02 | 0.98 |
| 3676-28 | 2424_E | 0.00 | 0.00 | 0.00 | 0.00 | 0.00 | 0.00 | 1.00 | 0.00 | 0.00 | 0.00 | 0.00 | 0.00 | F2-4 | **F2-4** | 0.36 | 0.64 |
| 3676-29 | 2424_H | 0.00 | 0.00 | 0.00 | 0.00 | 0.00 | 0.00 | 0.00 | 0.00 | 0.02 | 0.19 | 0.13 | 0.66 | Bput | ***B. puteoserpentis*** | 0.00 | 1.00 |
| 3676-30 | 3386_W | 0.01 | 0.56 | 0.00 | 0.00 | 0.00 | 0.00 | 0.43 | 0.00 | 0.00 | 0.00 | 0.00 | 0.00 | BC azo | **F2-4** | 0.54 | 0.46 |
| 3676-31 | 3386_X | 0.15 | 0.79 | 0.01 | 0.00 | 0.00 | 0.00 | 0.05 | 0.00 | 0.00 | 0.00 | 0.00 | 0.00 | BC azo | **F2-4** | 0.63 | 0.37 |
| 3676-32 | 3386_Y | 0.00 | 0.00 | 0.00 | 0.00 | 0.00 | 0.00 | 0.00 | 0.00 | 0.00 | 0.07 | 0.08 | 0.84 | Bput | ***B. puteoserpentis*** | 0.00 | 1.00 |
| 3676-33 | 3386_Z | 0.00 | 0.00 | 0.00 | 0.00 | 0.00 | 0.00 | 0.00 | 0.00 | 0.00 | 0.04 | 0.07 | 0.89 | Bput | ***B. puteoserpentis*** | 0.00 | 1.00 |
| 3676-37 | 3386_AA | 0.00 | 0.00 | 0.00 | 0.00 | 0.00 | 0.00 | 0.02 | 0.17 | 0.43 | 0.36 | 0.01 | 0.00 | BC put | **BC *puteoserpentis*** | 0.17 | 0.83 |
| 3676-38 | 3386_AB | 0.00 | 0.00 | 0.00 | 0.00 | 0.00 | 0.00 | 0.00 | 0.00 | 0.01 | 0.12 | 0.10 | 0.78 | Bput | ***B. puteoserpentis*** | 0.00 | 1.00 |
| 3125-1 | 3386_AC | 0.00 | 0.00 | 0.00 | 0.00 | 0.00 | 0.00 | 0.00 | 0.00 | 0.05 | 0.32 | 0.14 | 0.49 | Bput | ***B. puteoserpentis*** | 0.01 | 0.99 |
| 3125-2 | 3386_AD | 0.02 | 0.82 | 0.00 | 0.00 | 0.00 | 0.01 | 0.15 | 0.00 | 0.00 | 0.00 | 0.00 | 0.00 | BC azo | **F2-4** | 0.57 | 0.43 |
| 3125-3 | 3386_AE | 0.00 | 0.00 | 0.00 | 0.00 | 0.00 | 0.00 | 0.00 | 0.00 | 0.00 | 0.04 | 0.07 | 0.89 | Bput | ***B. puteoserpentis*** | 0.00 | 1.00 |
| 3125-4 | 3386_AF | 0.00 | 0.00 | 0.00 | 0.00 | 0.00 | 0.00 | 0.00 | 0.00 | 0.08 | 0.42 | 0.18 | 0.33 | BC put | **BC *puteoserpentis*** | 0.02 | 0.98 |
| 3125-5 | 3386_AG | 0.00 | 0.04 | 0.00 | 0.00 | 0.00 | 0.06 | 0.90 | 0.00 | 0.00 | 0.00 | 0.00 | 0.00 | F2-4 | **F2-4** | 0.43 | 0.57 |
| 3125-6 | 3386_AH | 0.00 | 0.32 | 0.00 | 0.00 | 0.00 | 0.00 | 0.68 | 0.00 | 0.00 | 0.00 | 0.00 | 0.00 | F2-4 | **F2-4** | 0.51 | 0.49 |
| 3125-7 | 3386_AI | 0.05 | 0.52 | 0.00 | 0.00 | 0.00 | 0.00 | 0.42 | 0.00 | 0.00 | 0.00 | 0.00 | 0.00 | BC azo | **F2-4** | 0.57 | 0.43 |
| 3125-8 | 2424_N | 0.00 | 0.03 | 0.00 | 0.00 | 0.00 | 0.00 | 0.96 | 0.00 | 0.00 | 0.00 | 0.00 | 0.00 | F2-4 | **F2-4** | 0.45 | 0.55 |
| 3125-9 | 3386_AJ | 0.00 | 0.00 | 0.00 | 0.00 | 0.00 | 0.00 | 1.00 | 0.00 | 0.00 | 0.00 | 0.00 | 0.00 | F2-4 | **F2-4** | 0.37 | 0.63 |
| 3125-10 | 3386_AK | 0.00 | 0.00 | 0.00 | 0.00 | 0.00 | 0.00 | 0.00 | 0.00 | 0.01 | 0.11 | 0.10 | 0.78 | Bput | ***B. puteoserpentis*** | 0.00 | 1.00 |
| 3125-11 | 3386_AL | 0.00 | 0.09 | 0.00 | 0.00 | 0.00 | 0.01 | 0.91 | 0.00 | 0.00 | 0.00 | 0.00 | 0.00 | F2-4 | **F2-4** | 0.45 | 0.55 |

**Supplementary Table S 3 | Statistics of SOX symbiont MAGs.** Complete: completeness based on gammaproteobacterial marker genes in CheckM; Contam (Strain): Contamination and percentage of this contamination which can be explained by strain heterogeneity according to CheckM; Contam Corr: Contamination after correction for strain heterogeneity (i.e. strain variants were not considered as contamination).

| MAG ID | Complete [%] | Contam (Strain) [%] | Contam Corr [%] | # Contigs | GC [%] | Genome size [Mb] | Read coverage [x] |
|---|---|---|---|---|---|---|---|
| SOX_BS_2424_D | 94.25 | 0 (0) | 0.00 | 914 | 0.37 | 2.40 | 22 |
| SOX_BS_2424_E | 93.69 | 0.16 (33.33) | 0.11 | 3206 | 0.37 | 2.89 | 100 |
| SOX_BS_2424_H | 92.28 | 0 (0) | 0.00 | 772 | 0.37 | 2.13 | 17 |
| SOX_BS_2424_N | 92.28 | 6.87 (95.45) | 0.31 | 4717 | 0.36 | 3.57 | 1001 |
| SOX_BS_3386_A | 94.44 | 1.11 (66.67) | 0.37 | 3998 | 0.37 | 2.76 | 241 |
| SOX_BS_3386_AB | 94.25 | 1.64 (100) | 0.00 | 4095 | 0.37 | 2.80 | 372 |
| SOX_BS_3386_AD | 94.25 | 0.14 (100) | 0.00 | 3928 | 0.37 | 2.69 | 472 |
| SOX_BS_3386_AE | 94.25 | 0.28 (100) | 0.00 | 3899 | 0.37 | 2.73 | 105 |
| SOX_BS_3386_AG | 95.09 | 7.24 (84) | 1.16 | 4767 | 0.36 | 3.46 | 1744 |
| SOX_BS_3386_AH | 94.25 | 3.5 (12.5) | 3.06 | 4113 | 0.36 | 3.44 | 542 |
| SOX_BS_3386_AJ | 92 | 8.81 (86.11) | 1.22 | 5570 | 0.36 | 4.04 | 989 |
| SOX_BS_3386_AK | 94.63 | 0.45 (40) | 0.27 | 3648 | 0.37 | 2.70 | 301 |
| SOX_BS_3386_AL | 94.25 | 3.04 (58.33) | 1.27 | 3415 | 0.37 | 2.62 | 387 |
| SOX_BS_3386_C | 94.25 | 3.51 (8.33) | 3.22 | 3038 | 0.37 | 2.50 | 502 |
| SOX_BS_3386_D | 94.25 | 2.09 (25) | 1.57 | 4408 | 0.37 | 2.92 | 432 |
| SOX_BS_3386_E | 90.18 | 0 (0) | 0.00 | 143 | 0.38 | 1.43 | 488 |
| SOX_BS_3386_F | 94.25 | 2.2 (100) | 0.00 | 4006 | 0.37 | 2.70 | 382 |
| SOX_BS_3386_G | 94.25 | 2.57 (90.91) | 0.23 | 3656 | 0.37 | 2.63 | 601 |
| SOX_BS_3386_H | 89.05 | 0 (0) | 0.00 | 145 | 0.38 | 1.37 | 527 |
| SOX_BS_3386_I | 92.43 | 0 (0) | 0.00 | 157 | 0.38 | 1.48 | 365 |
| SOX_BS_3386_J | 91.3 | 0 (0) | 0.00 | 163 | 0.38 | 1.48 | 2588 |
| SOX_BS_3386_M | 94.25 | 4.45 (33.33) | 2.97 | 3425 | 0.37 | 2.70 | 2704 |
| SOX_BS_3386_O | 94.81 | 3.79 (36.36) | 2.41 | 4500 | 0.37 | 3.07 | 515 |
| SOX_BS_3386_S | 94.25 | 4.03 (69.23) | 1.24 | 3607 | 0.37 | 2.90 | 489 |

| | | | | | | | |
|---|---|---|---|---|---|---|---|
| SOX_BS_3386_T | 93.97 | 4.35 (63.64) | 1.58 | 2934 | 0.37 | 2.60 | 298 |
| SOX_BS_3386_U | 94.25 | 0.14 (100) | 0.00 | 4081 | 0.37 | 2.81 | 249 |
| SOX_BS_3386_W | 94.25 | 2.2 (87.5) | 0.28 | 3531 | 0.37 | 2.72 | 398 |
| SOX_BS_3386_X | 94.25 | 4.57 (28.57) | 3.21 | 3161 | 0.37 | 2.79 | 419 |
| SOX_BS_3386_Y | 94.25 | 0.14 (100) | 0.00 | 3032 | 0.37 | 2.66 | 431 |
| SOX_BS_3386_Z | 94.53 | 3.31 (61.54) | 1.27 | 4674 | 0.36 | 3.48 | 951 |

**Supplementary Table S 4 | Overview of analyses of Bathymodiolus SOX symbionts.**

| Analysis | Input data/reference | Level of resolution |
|---|---|---|
| Phylogenomics | 171 marker genes extracted from NMAR MAGs | symbiont subspecies |
| Average nucleotide identity | Broken Spur MAGs | symbiont subspecies |
| Gene abundances | Mussel metagenomes from Broken Spur mapped against Broken Spur orthologous gene catalogue | symbiont strain |
| SNP analyses | Mussel metagenomes from Broken Spur mapped against Broken Spur orthologous gene catalogue | symbiont strain |
| Redundancy analysis | Mussel metagenomes from NMAR sites mapped against NMAR orthologous gene catalogue | symbiont strain |

**Supplementary Table S 5 | List of external data.** BioProject and dataset accessions for the European Nucleotide Archive and the respective publication were listed if available. Lat: Latitude; Lon: Longitude; Complete: completeness based on gammaproteobacterial marker genes in CheckM; Contam: Contamination; Strain: Percentage of this contamination which can be explained by strain heterogeneity according to CheckM; Pub: Publication (* = Ansorge et al., in prep.).

| MAG | Site | Lat | Lon | Cruise | Host species | Complete | Contam | Strain | Pub | BioProject | Accession |
|---|---|---|---|---|---|---|---|---|---|---|---|
| 1048F | Lucky Strike (Montsegur) | 37.288 | -32.276 | Biobaz (2013) | *B. azoricus* | 94.53 | 0 | 0 | * | PRJEB36091 | GCA_903813355 |
| 1048G | Lucky Strike (Montsegur) | 37.288 | -32.276 | Biobaz (2013) | *B. azoricus* | 94.53 | 0 | 0 | * | PRJEB36091 | GCA_903813365 |
| 1048H | Lucky Strike (Montsegur) | 37.288 | -32.276 | Biobaz (2013) | *B. azoricus* | 95.09 | 0 | 0 | * | PRJEB36091 | GCA_903819415 |
| 1048I | Lucky Strike (Eiffel Tower) | 37.283 | -32.276 | Biobaz (2013) | *B. azoricus* | 93.97 | 0.03 | 0 | * | PRJEB36091 | GCA_903819345 |
| 1048J | Lucky Strike (Eiffel Tower) | 38.283 | -32.276 | Biobaz (2013) | *B. azoricus* | 94.53 | 0.56 | 100 | * | PRJEB36091 | GCA_903819365 |
| 1586B | Lucky Strike (Eiffel Tower) | 37.289 | -32.275 | Biobaz (2013) | *B. azoricus* | 93.41 | 0.84 | 100 | * | PRJEB36091 | GCA_903813405 |
| 1586C | Lucky Strike (Eiffel Tower) | 37.289 | -32.275 | Biobaz (2013) | *B. azoricus* | 92.85 | 0.84 | 50 | * | PRJEB36091 | GCA_903819425 |
| 1586D | Lucky Strike (Eiffel Tower) | 37.289 | -32.275 | Biobaz (2013) | *B. azoricus* | 93.97 | 0 | 0 | * | PRJEB36091 | GCA_903813415 |
| 1586E | Lucky Strike (Eiffel Tower) | 37.289 | -32.275 | Biobaz (2013) | *B. azoricus* | 92.85 | 0 | 0 | * | PRJEB36091 | GCA_903813425 |
| 1586F | Lucky Strike (Eiffel Tower) | 37.283 | -32.276 | Biobaz (2013) | *B. azoricus* | 92.85 | 1.4 | 33.3 | * | PRJEB36091 | GCA_903813665 |

| | | | | | | | | | | | |
|---|---|---|---|---|---|---|---|---|---|---|---|
| 1586G | Lucky Strike (Eiffel Tower) | 37.283 | -32.276 | Biobaz (2013) | *B. azoricus* | 93.97 | 0 | 0 | * | PRJEB36091 | GCA_903813675 |
| 1586I | Lucky Strike (Eiffel Tower) | 37.283 | -32.276 | Biobaz (2013) | *B. azoricus* | 94.53 | 0 | 0 | | | SAMEA6822959 |
| 1586J | Lucky Strike (Eiffel Tower) | 37.283 | -32.276 | Biobaz (2013) | *B. azoricus* | 93.97 | 0.56 | 0 | * | PRJEB36091 | GCA_903799955 |
| 1586K | Lucky Strike (Montsegur) | 37.288 | -32.276 | Biobaz (2013) | *B. azoricus* | 92.28 | 0.56 | 0 | * | PRJEB36091 | GCA_903813645 |
| 1586N | Lucky Strike (Montsegur) | 37.288 | -32.276 | Biobaz (2013) | *B. azoricus* | 94.53 | 0 | 0 | * | PRJEB36091 | GCA_903813615 |
| 1586O | Lucky Strike (Montsegur) | 37.288 | -32.276 | Biobaz (2013) | *B. azoricus* | 93.6 | 1.12 | 50 | * | PRJEB36091 | GCA_903813655 |
| 1586P | Menez Gwen (White Flames) | 37.844 | -31.519 | Biobaz (2013) | *B. azoricus* | 91.72 | 0 | 0 | * | PRJEB36091 | GCA_903813625 |
| 1586Q | Menez Gwen (White Flames) | 37.844 | -31.519 | Biobaz (2013) | *B. azoricus* | 92.28 | 0 | 0 | * | PRJEB36091 | GCA_903813695 |
| 1586R | Menez Gwen (White Flames) | 37.844 | -31.519 | Biobaz (2013) | *B. azoricus* | 93.97 | 0 | 0 | | | SAMEA6822960 |
| 1586S | Menez Gwen (White Flames) | 37.844 | -31.519 | Biobaz (2013) | *B. azoricus* | 93.97 | 0 | 0 | * | PRJEB36091 | GCA_903813685 |
| 1600F | Rainbow | 36.229 | -33.902 | Biobaz (2013) | *B. azoricus* | 93.97 | 0 | 0 | * | PRJEB36091 | GCA_903813635 |
| 1600G | Rainbow | 36.229 | -33.902 | Biobaz (2013) | *B. azoricus* | 93.97 | 0 | 0 | * | PRJEB36091 | GCA_903813705 |
| 1600H | Rainbow | 36.229 | -33.902 | Biobaz (2013) | *B. azoricus* | 93.93 | 0 | 0 | * | PRJEB36091 | GCA_903819385 |

| | | | | | | | | | | | |
|---|---|---|---|---|---|---|---|---|---|---|---|
| 1600I | Rainbow | 36.229 | -33.902 | Biobaz (2013) | *B. azoricus* | 91.72 | 0 | 0 | * | PRJEB36091 | GCA_903813715 |
| 1600J | Rainbow | 36.229 | -33.902 | Biobaz (2013) | *B. azoricus* | 93.97 | 0.14 | 100 | * | PRJEB36091 | GCA_903813725 |
| BBROOKSOX | Chapopote (Mexico) | 21.90002 | -93.4353 | M114-2 | *B. brooksi* | 93.27 | 0.56 | 0 | | PRJEB17996 | GCA_900128405.1 |
| BHECKSOX | Chapopote (Mexico) | 21.90005 | -93.4354 | M114-2 | *B. heckerae* | 92.66 | 4.31 | 63.6 | | PRJEB17996 | GCA_900128515.1 |
| 1115A | Semenov | 13.513 | -44.963 | Odemar (2014) | *B. puteoserpentis* | 94.53 | 3.09 | 75 | * | PRJEB36091 | GCA_903813375 |
| 1115B | Semenov | 13.513 | -44.963 | Odemar (2014) | *B. puteoserpentis* | 93.41 | 3.09 | 85.7 | * | PRJEB36091 | GCA_903813385 |
| 1115C | Semenov | 13.513 | -44.963 | Odemar (2014) | *B. puteoserpentis* | 94.53 | 0.56 | 100 | * | PRJEB36091 | GCA_903813395 |
| 2065A | Logatchev Quest | 14.753 | -44.979 | M64-2 Logatchev (2005) | *B. puteoserpentis* | 94.72 | 7.3 | 100 | * | PRJEB36091 | GCA_903813905 |
| 2065B | Logatchev Quest | 14.753 | -44.979 | M64-2 Logatchev (2005) | *B. puteoserpentis* | 94.72 | 1.15 | 100 | * | PRJEB36091 | GCA_903813925 |
| 2487A | Logatchev Quest | 14.753 | -44.980 | M126 (2016) | *B. puteoserpentis* | 94.16 | 1.4 | 57.1 | * | PRJEB36091 | GCA_903819405 |
| 2487B | Logatchev Quest | 14.753 | -44.980 | M126 (2016) | *B. puteoserpentis* | 94.72 | 1.4 | 57.1 | * | PRJEB36091 | GCA_903819245 |
| 2487C | Logatchev Quest | 14.753 | -44.980 | M126 (2016) | *B. puteoserpentis* | 94.72 | 1.4 | 83.3 | * | PRJEB36091 | GCA_903813875 |
| 2487D | Semenov | 13.514 | -44.963 | M126 (2016) | *B. puteoserpentis* | 94.53 | 1.12 | 66.7 | * | PRJEB36091 | GCA_903813855 |
| 2487E | Semenov | 13.514 | -44.963 | M126 (2016) | *B. puteoserpentis* | 94.72 | 1.12 | 66.7 | * | PRJEB36091 | GCA_903819375 |
| 2487F | Semenov | 13.514 | -44.963 | M126 (2016) | *B. puteoserpentis* | 94.53 | 1.97 | 83.3 | * | PRJEB36091 | GCA_903813935 |

| | | | | | | | | | | | |
|---|---|---|---|---|---|---|---|---|---|---|---|
| 3722CJ | Logatchev Quest | 14.753 | -44.981 | MSM10-03 Hydromar VII | *B. puteoserpentis* | 94.72 | 5.15 | 50 | * | PRJEB36091 | GCA_903814125 |
| 3722CK | Logatchev Quest | 14.753 | -44.981 | MSM10-03 Hydromar VII | *B. puteoserpentis* | 94.72 | 3.65 | 78.6 | * | PRJEB36091 | GCA_903819395 |
| 3722CL | Logatchev Quest | 14.753 | -44.981 | MSM10-03 Hydromar VII | *B. puteoserpentis* | 94.72 | 3.93 | 92.9 | * | PRJEB36091 | GCA_903814105 |
| 3722CM | Logatchev Quest | 14.753 | -44.981 | MSM10-03 Hydromar VII | *B. puteoserpentis* | 94.72 | 3.23 | 100 | * | PRJEB36091 | GCA_903814155 |
| 3722CN | Logatchev Quest | 14.753 | -44.981 | MSM10-03 Hydromar VII | *B. puteoserpentis* | 93.6 | 2.39 | 76.9 | * | PRJEB36091 | GCA_903814145 |
| 3722CO | Logatchev Quest | 14.753 | -44.981 | MSM10-03 Hydromar VII | *B. puteoserpentis* | 93.03 | 2.81 | 92.3 | * | PRJEB36091 | GCA_903814115 |
| 3722CP | Logatchev Quest | 14.753 | -44.980 | M126 (2016) | *B. puteoserpentis* | 94.72 | 8.8 | 90.9 | * | PRJEB36091 | GCA_903814135 |
| Endosymbiont of *Bathymodiolus septemdierum* str. Myojin knoll DNA, complete genome | Izu-Bonin Arc, Myojin knoll (Japan) | 32.104 | 139.219 | | *B. septemdierum* | 94.83 | 0.56 | 100 | (Ikuta et al., 2016) | PRJDB949 | NZ_AP013042.1 |

| | | | | | | | | | | | |
|---|---|---|---|---|---|---|---|---|---|---|---|
| C112 | Clueless | -4.803 | -12.372 | M78-2 (2009) | *B.* sp. Clueless | 94.53 | 1.31 | 75 | * | PRJEB36091 | GCA_903814195 |
| C113 | Clueless | -4.803 | -12.372 | M78-2 (2009) | *B.* sp. Clueless | 94.53 | 1.4 | 66.7 | * | PRJEB36091 | GCA_903814185 |
| C114 | Clueless | -4.803 | -12.372 | M78-2 (2009) | *B.* sp. Clueless | 94.53 | 1.4 | 66.7 | * | PRJEB36091 | GCA_903814175 |
| L102 | Lilliput | -9.547 | -13.210 | M78-2 (2009) | *B.* sp. Lilliput | 93.33 | 1.12 | 100 | * | PRJEB36091 | GCA_903813445 |
| L51 | Lilliput | -9.547 | -13.210 | M78-2 (2009) | *B.* sp. Lilliput | 94.08 | 1.69 | 100 | * | PRJEB36091 | GCA_903813485 |
| L54 | Lilliput | -9.547 | -13.210 | M78-2 (2009) | *B.* sp. Lilliput | 94.36 | 0.84 | 100 | * | PRJEB36091 | GCA_903813455 |
| *Bathymodiolus thermophilus* thioautotrophic gill symbiont strain:BAT/CrabSpa'14 | East Pacific Rise (EPR) 9°N | 9.839833 | -104.292 | R/V *Atlantis* cruise AT26–10 | *B. thermophilus* | 96.98 | 11.32 | 81.4 | (Ponnudurai et al., 2017) | PRJNA339702 | GCA_001875585 |
| *Ca.* Ruthia magnifica str. Cm | 9° East Pacific Rise vent field | 9.830 | -104.290 | | *C. magnifica* | 86.67 | 0 | 0 | (Newton et al., 2007) | PRJNA16841 | CP000488 |
| *Ca.* Vesicomyosocius okutanii HA | Sagami Bay | 35.117 | 139.383 | | *C. okutanii* | 85.69 | 0 | 0 | (Kuwahara et al., 2007) | PRJDA18267 | AP009247 |
| *Ca.* Thioglobus autotrophicus strain EF1 | Effingham Inlet (estimated coordinates) | 49.029 | -125.154 | | | 94.64 | 0 | 0 | (Shah & Morris, 2015) | PRJNA224116 | NZ_CP010552 |
| *Ca.* Thioglobus singularis PS1 | Puget Sound | 47.600 | -122.450 | | | 94.36 | 0 | 0 | (Marshall & Morris, 2015) | PRJNA229178 | CP006911 |

| | | | | | | | | | | | |
|---|---|---|---|---|---|---|---|---|---|---|---|
| *Thiomicrospira crunogena* XCL-2 (*Hydrogenovibrio crunogenus* XCL-2) | | | | | | 99.72 | 0.19 | 0 | (Scott et al., 2006) | PRJNA13018 | NC_007520.2 |
